# *Chlamydia trachomatis* restricts signaling through NOD2 until late in the pathogen’s developmental cycle

**DOI:** 10.1101/2025.05.28.656691

**Authors:** Grace Overman, Iris Loeckener, Zachary Williford, Sung Davis, Aissata Diallo, Josie Blair, Beate Henrichfreise, George W. Liechti

## Abstract

Pathogenic chlamydial species restrict their peptidoglycan (PG) to the division septum of their replicative forms. PG is a microbe-associated molecular pattern (MAMP) and two of its major pattern recognition receptors in human cells are nucleotide-binding oligomerization domain-containing proteins 1 and 2 (NOD1 and NOD2, respectively). It has been proposed that this unique morphological feature is evidence of pathoadaptation by the microbe, permitting PG-dependent cell division while also reducing the bacterium’s recognition by innate immune receptors. *Chlamydia trachomatis*-infected cells activate NOD1 signaling within 8-12 hours of exposure to the bacterium, roughly coinciding with the microbe’s transition from its infectious to replicative forms. Here we report that, unlike NOD1 signaling, *Chlamydia*-induced NOD2 signaling does not occur until later in the pathogen’s developmental cycle. Both *C. trachomatis* and the related murine pathogen *Chlamydia muridarum* signal late in infection in HEK293 reporter cell lines expressing either human or murine-derived NOD2 receptors. NOD2 signaling can be modulated by disruption of the chlamydial amidase enzyme, AmiA_CT_, interrupting the microbe’s developmental cycle, and treatment with inhibitors of lipooligosaccharide or peptidoglycan biosynthesis / assembly. These results mirror prior observations with Chlamydia-induced TLR9 signaling, leading us to hypothesize that Chlamydia-induced NOD2 signaling results from lytic events that occur sporadically during the transition between the pathogen’s developmental forms. Given our finding that pre-treating cells with NOD2-stimulatory ligands reduces chlamydial inclusion size and delays the developmental cycle, we hypothesize that the microbe preferentially degrades its PG during development to reduce the generation of NOD2 ligands.

## INTRODUCTION

*Chlamydia trachomatis* is the leading cause of bacterial sexually transmitted disease and infectious blindness in the world, with over 100 million estimated cases of sexually transmitted infection^1^ and ∼100 million people still at risk of trachoma-induced blindness^2^. This obligate, intracellular microbe has evolved a number of pathoadaptive means of circumventing the human immune system at the cellular level. Though this pathogen is susceptible to many commercial antibiotics, the estimated burden within the global population remains high due to the large majority of infections being asymptomatic^3,4^, enabling the organism to spread in the absence of effective screening and treatment.

One of the adaptations made by chlamydial species to life inside of host organisms is the utilization of a biphasic developmental cycle^5^. Infectious forms, termed Elementary bodies (EBs) have reduced metabolic rates^6^ and utilize lipooligosaccharide and gDNA with significantly reduced immunostimulatory properties^7–11^. Upon entering into host cells, chlamydial EBs transition into Reticulate bodies (RBs), and begin replicating via a unique cell division process^12^ that is dependent on the synthesis and turnover of peptidoglycan (PG)^13–15^. In most other bacterial species, to include some chlamydial-related organisms^16^, PG is a major component of the bacterial cell wall (termed a sacculus) that confers strength and rigidity to bacterial cells, while also allowing microbes to maintain or alter their overall shape^17–19^. Pathogenic chlamydial species lack sacculi, but synthesize PG at their division planes^13,15,20^. Paradoxically, because they lack FtsZ^21^, the major organizer of septal PG in almost all bacterial species^22^, they utilize the PG synthesis machinery primarily associated with sacculus construction in other bacterial species^15,23–29^. While encoding two PG synthase complexes primarily associated with either side-wall^30^ or septal^31^ PG biosynthesis, *C. trachomatis* utilizes both to modulate the dimensions of its ‘PG ring’ during the initial and terminal stages of its division process^32,33^.

It has been proposed that the absence of PG sacculi in chlamydial species is evidence of pathoadaptation by these organisms^13,15,34,35^, due to PG being a major immunostimulatory molecule that is recognized by a wide range of host cellular receptors^36^. As PG is only present in the organism’s replicative forms and in relatively low abundance when compared to the amount required to generate sacculi, it stands to reason that reducing the abundance of PG-derived muropeptides would directly impact *Chlamydia*-host cell interaction(s). This hypothesis is supported in part by the characterization of a rudimentary PG-recycling pathway encoded by *C. trachomatis* that enables the reabsorption of meso-diaminopimelic acid (mDAP)– containing muropeptides^37,38^, further reducing its immunostimulatory profile.

The two most well-studied immunoreceptors for bacterial PG recognition are the NOD-like receptors (NLRs) NOD1 and NOD2^39^. NOD1 recognizes PG-derived peptides containing mDAP^40^ while NOD2 primarily recognizes peptides containing N-acetylmuramic acid (MurNAc), the principle component of NOD2 / CARD15 stimulatory ligands^41^. While it has been over a decade since PG was first demonstrated to be present in these organisms^13^, the degree to which these receptors play a role during Chlamydia infections is still in question^42^. Correlative studies examining patient cohorts have noted an association between NOD1 functionality and bacterial clearance / disease pathology^43,44^ while a less robust association has been observed between reduced NOD2 functionality and an increased risk of tubal pathology subsequent to chlamydial infection^45^. Both receptors are expressed throughout the female reproductive tract^46,47^, with some evidence that expression of NOD2 (but not NOD1) is impacted by phases of the menstrual cycle^48^. NOD1 induction has been demonstrated to have a significant role in inducing inflammation in vitro^49–51^, and murine studies have linked both receptors to Endoplasmic Reticulum Stress-Induced Inflammation during chlamydial infection and NOD1/2-deficient mice exhibit a reduction in clearance rates^52,53^, though it is presently unclear if these phenotypes are associated with ligand-dependent or ligand-independent NOD1/2 signaling^54^.

We have previously reported that PG can be visualized in cells infected with *C. trachomatis* as early as 8 hours post-infection (hpi)^13^, coinciding with the transition of EBs to RBs and transcriptome data demonstrating the expression of chlamydial genes associated with PG biosynthesis and turnover^55^. NOD1 signaling can be observed within 24 hpi in cells infected with *C. trachomatis*, and we have previously demonstrated that treatment with various antibiotics, metal chelators, and host cell cytokines can impact the intensity of chlamydia-induced NOD1 signaling^56^. Interestingly, while NOD2- stimulatory ligands have been reported to be present in lysates generated from Chlamydia-infected cells^57^, no signaling was observed 24 hpi when reporter cells were infected directly, as noted by Packiam (personal communication, April 2014). Given our recent work demonstrating that TLR9 signaling is delayed in Chlamydia-infected cells and appears to be dependent on either bacterial lysis or transitioning between chlamydial forms^9^, we reasoned that this could potentially explain the discrepancy under these two conditions. We set out to establish the dynamics of Chlamydia-induced NOD2 signaling, and determine what factor(s) impact its amplitude and temporal kinetics.

## RESULTS

### *Chlamydia trachomatis* signals via NOD2 late in the pathogen’s developmental cycle

We utilized a secreted embryonic alkaline phosphatase (SEAP) reporter system to compare the relative signaling of human (h)NOD1 and hNOD2 in cells infected with *C. trachomatis*. HEK-Blue cells stably express hNOD1 and hNOD2 and an NFκB- inducible SEAP reporter gene. By comparing SEAP levels present in cell supernatants between hNOD-expressing cells and cells lacking hNOD expression (Null1 and Null2 cells), NOD-specific signaling can be assessed between controls and a variety of experimental conditions. SEAP was detected in the supernatants of *C. trachomatis* infected cells at elevated levels at 24 and 44 hours post infection (hpi) for NOD1 (**Fig. 1A**), however, NOD2 signaling appeared to be delayed as SEAP was only present at appreciable levels in cell supernatants at 44 hpi (**Fig. 1B**). In order to more closely follow the kinetics of *C. trachomatis*-induced NOD2 signaling, we conducted real-time monitoring of SEAP levels present in cell supernatants, beginning at 18hpi. Infected cells were incubated in HEK-Blue Detection cell culture medium, allowing hydrolysis of substrate by SEAP to be measured via microplate reader every 10 minutes for 28 hours. We found that SEAP begins to accumulate at detectable levels ∼19-20 hpi and that activity began to accelerate in a MOI-dependent manner at ∼24 hpi (MOI ∼2) and ∼30 hpi (MOI ∼0.2) (**Fig. 1C**). As these observations roughly overlap with the pathogen’s transition from its replicative to infectious form, we concluded that NOD2 ligand release may be the result of *C. trachomatis* undergoing developmental form conversion.

**Figure 1.**
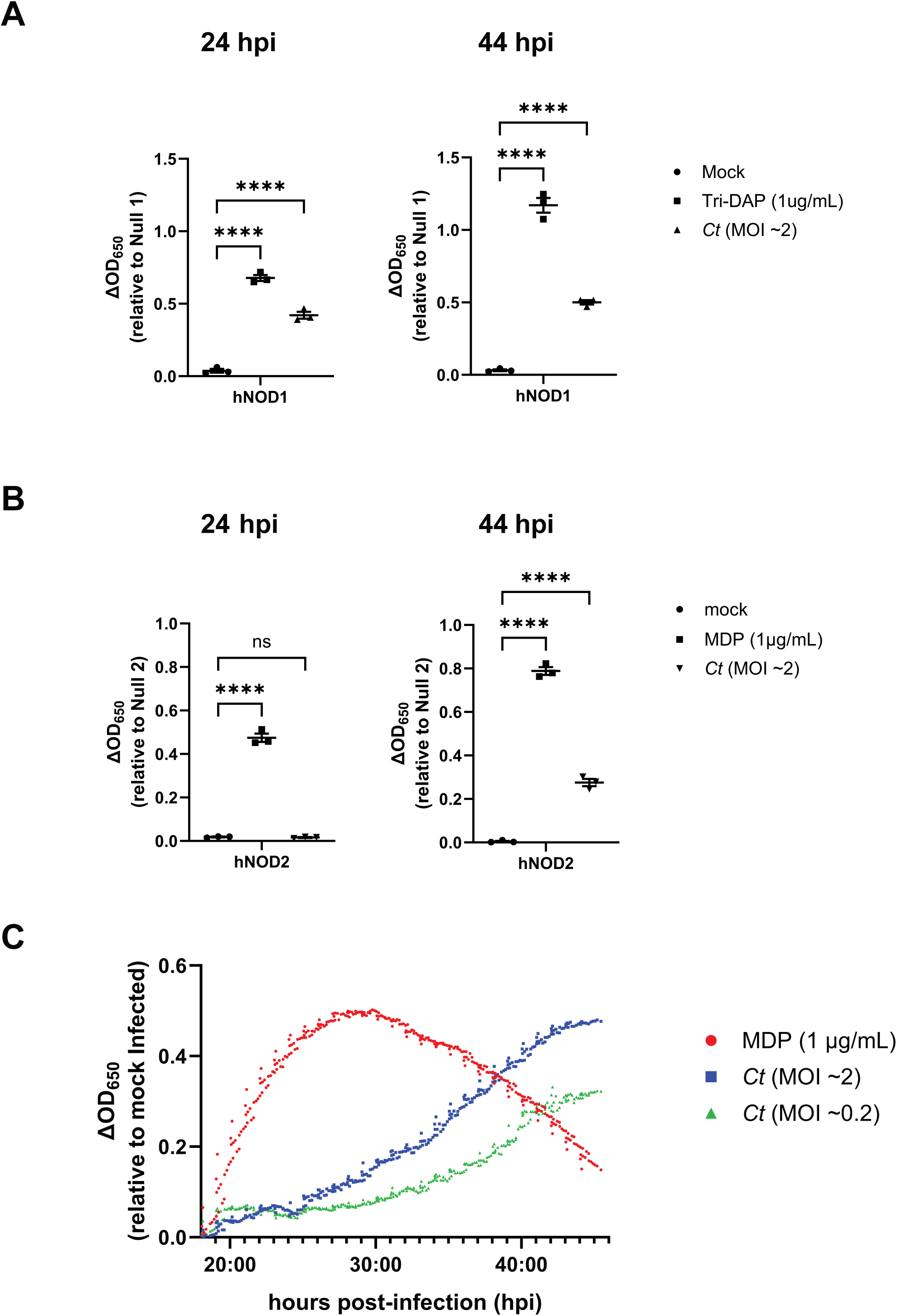
*Chlamydia trachomatis* signals via NOD2 late in the pathogen’s developmental cycle. SEAP activity was measured from the supernatants of hNOD1 **(A)** and hNOD2 **(B)** and Null1/2 HEK 293 reporter cells infected with *C. trachomatis* serovar L2 (strain Bu/434) at 24 and 44 hpi. Data points represent separate, biological replicates, with lines delineating the mean and error bars representing the standard error of the mean. Groups were compared via one-way ANOVA with multiple comparisons. ****; p ≤ 0.0001, ns; not significant. **(C)** SEAP activity was measured in the supernatants of infected NOD2-expressing HEK 293 reporter cells grown in HEK- Blue™ Detection media. OD_650_ values for each well were assessed every ten minutes beginning at 18 hpi, and values plotted are relative to mock infected control wells on the same plate. Data is representative of three wells per condition on the same plate (technical replicates). Positive control (muramyl dipeptide; MDP) was added to control wells at 18 hpi.

### Chlamydia-specific NOD2 signaling in HEK293 cells occurs as a result of developmental form conversion

We have previously reported that recognition of chlamydial species by TLR9 also occurs late in infection, and we proposed that chlamydial DNA is likely released due to random lytic events that occur during the transition from RBs to EBs^9^. To test whether interrupting the developmental cycle could prevent / dampen NOD2-stimulatory muropeptide release, we infected hNOD2 cells with *C. trachomatis*, and added chloramphenicol at various time points post infection to interrupt protein synthesis and subsequently prevent developmental form conversion. When protein synthesis was inhibited at 18 hpi, a time point prior to developmental form conversion^58^, NOD2 signaling decreased, and slowly increased when chloramphenicol was added at later and later time points (**Fig. 2A**). By contrast, when chlamydial LOS biosynthesis was inhibited with the use of a LpxC inhibitor (LPC-011^59,60^), NOD2 signaling was significantly increased when higher MOIs were used **(Fig. 2B)**. This enhancement did not appear to significantly impact the timing of NOD2-ligand release, as signaling was only observed at the 44hpi time point and only when higher MOIs were used. These observations, in addition to the kinetics of Chlamydia-specific NOD2 signaling **(Fig 1C)**, support the premise that NOD2 ligand release from *C. trachomatis* is likely a result of developmental form conversion.

**Figure 2.**
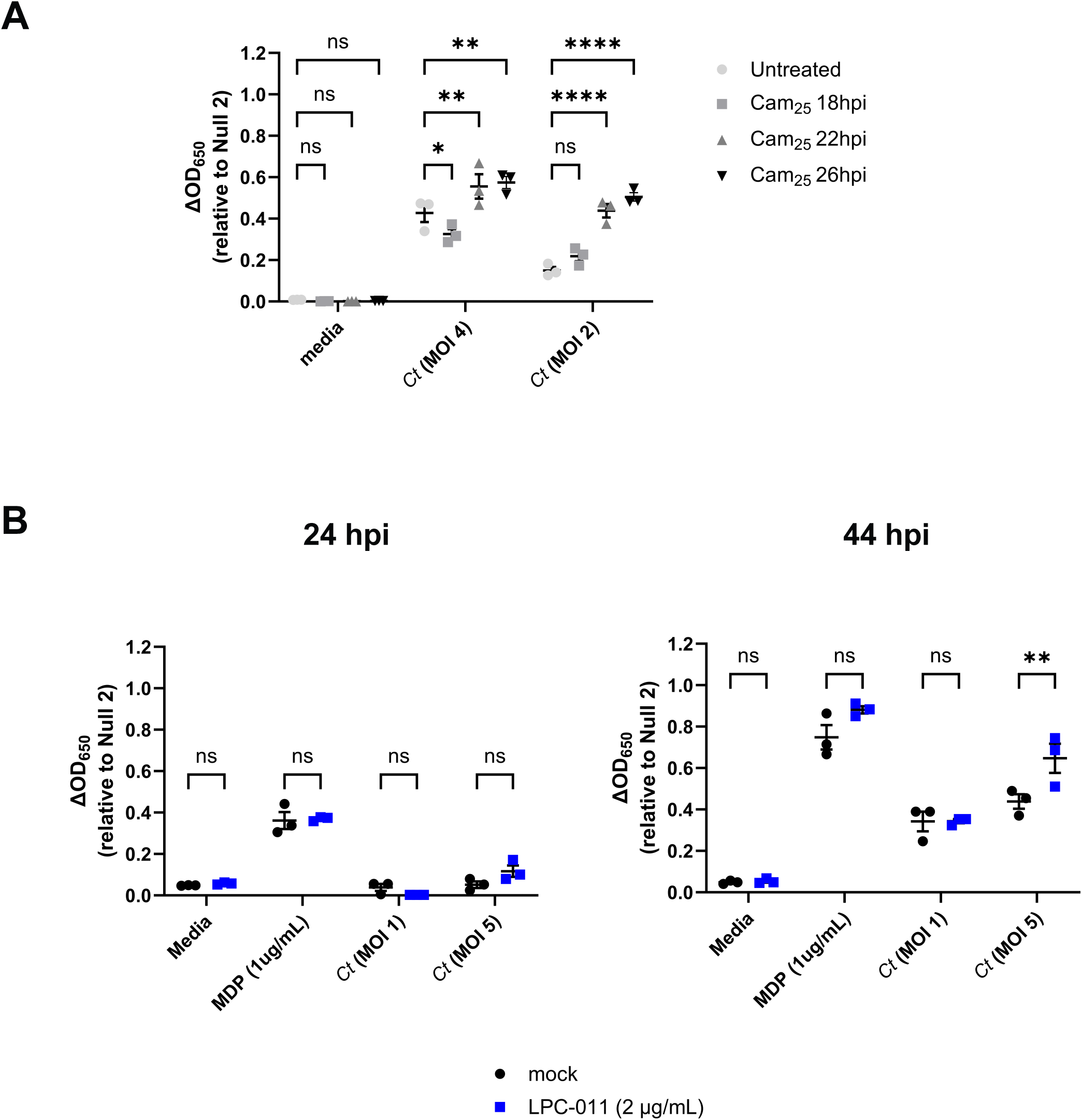
Chlamydia-specific NOD2 signaling in HEK293 cells occurs as a result of developmental form conversion. **(A)** *C. trachomatis*-infected hNOD2-HEK293 reporter cells were treated with chloramphenicol (25 ug/mL; Cam_25_) at the time points indicated and SEAP activity was measured at 44hpi. **(B,C)** The effects of the LOS inhibitor LPC-011 on *C. trachomatis*-induced NOD2 signaling at 24 **(B)** and 44 **(C)** hpi. Data points represent three independent, biological replicates and error bars represent standard error of the mean. Groups were compared via two-way ANOVA with multiple comparisons. ****; p ≤ 0.0001, **; p≤ 0.01, *; p ≤ 0.05, ns, not significant.

### Knock-down of *amiA_Ct_* impacts NOD2 signaling

Assuming that bacterial lysis is the cause for the release of NOD2-stimulatory ligands in Chlamydia-infected cells, it is not readily apparent why the release of Chlamydia-specific, NOD1-stimulatory ligands is not similarly impacted. As mentioned previously, *C. trachomatis* encodes a rudimentary peptidoglycan recycling pathway that enables it to limit the amount of meso-DAP-containing peptidoglycan fragment release into the intracellular environment^37,38^, however, to date no pathway for the recycling of anhydroMurNAc has been characterized in any Chlamydia species.

We questioned whether the molecular process(es) of PG degradation employed by the pathogen potentially influence its NLR stimulatory potential. *C. trachomatis* and other *Chlamydia spp*. and chlamydia-related organisms make use of two separate degradation enzymes in order to remodel their septal PG during the initiation and termination of their division process: an amidase (AmiA) that cleaves the stem peptide from MurNAc^61^ and a lytic transglycosylase (SpoIID) that cleaves denuded glycan strands^62^. MurNAc alone is a poor stimulator of NOD2 receptors, and most stimulatory ligands require at least two amino acids of the PG stem peptide to be present, in the correct spatial isomerism, in order for MurNAc to bind sufficiently well to result in a signaling cascade^63^. We reasoned that under normal conditions inherent to the PG turnover found in *Chlamydia* species, sufficient amidase activity would effectively eliminate any and all amino acids attached to MurNAc, thus fundamentally skewing NLR signaling towards NOD1 (which is mDAP-specific^40^) and away from NOD2.

In order to assess the impact of the chlamydial amidase on NOD2 signaling, we utilized a previously characterized CRISPRi-knockdown approach^64^ in order to gauge how reducing amidase activity would impact NOD signaling in *C. trachomatis*-infected cells. *C. trachomatis* strain -pL2 was transformed with either an empty plasmid; pLCria (NT), a knockdown construct; pLCria (*amiA*), or a knockdown construct with a HIS-tagged complementation allele; pLCria-amiA_6xHIS (amiA) (Dannenberg *et al*., in prep).

Each strain was then titered and used to infect hNOD2-expressing HEK293 reporter cells under either native or inducing conditions. Counter to our initial hypothesis, the transformant containing the AmiA knockdown plasmid appeared to exhibit slightly enhanced baseline NOD2 signaling when compared to control strains, a trend that reached statistical significance when higher MOIs were used (**Fig. 3A, left column**). When the three strains were compared using the lower MOI, the AmiA knockdown strain exhibited a significant (p ≤ 0.0001) decrease in NOD2 signaling (**Fig. 3B, right column**), a result that did not reach significance in either empty vector or complementation control strains.

**Figure 3.**
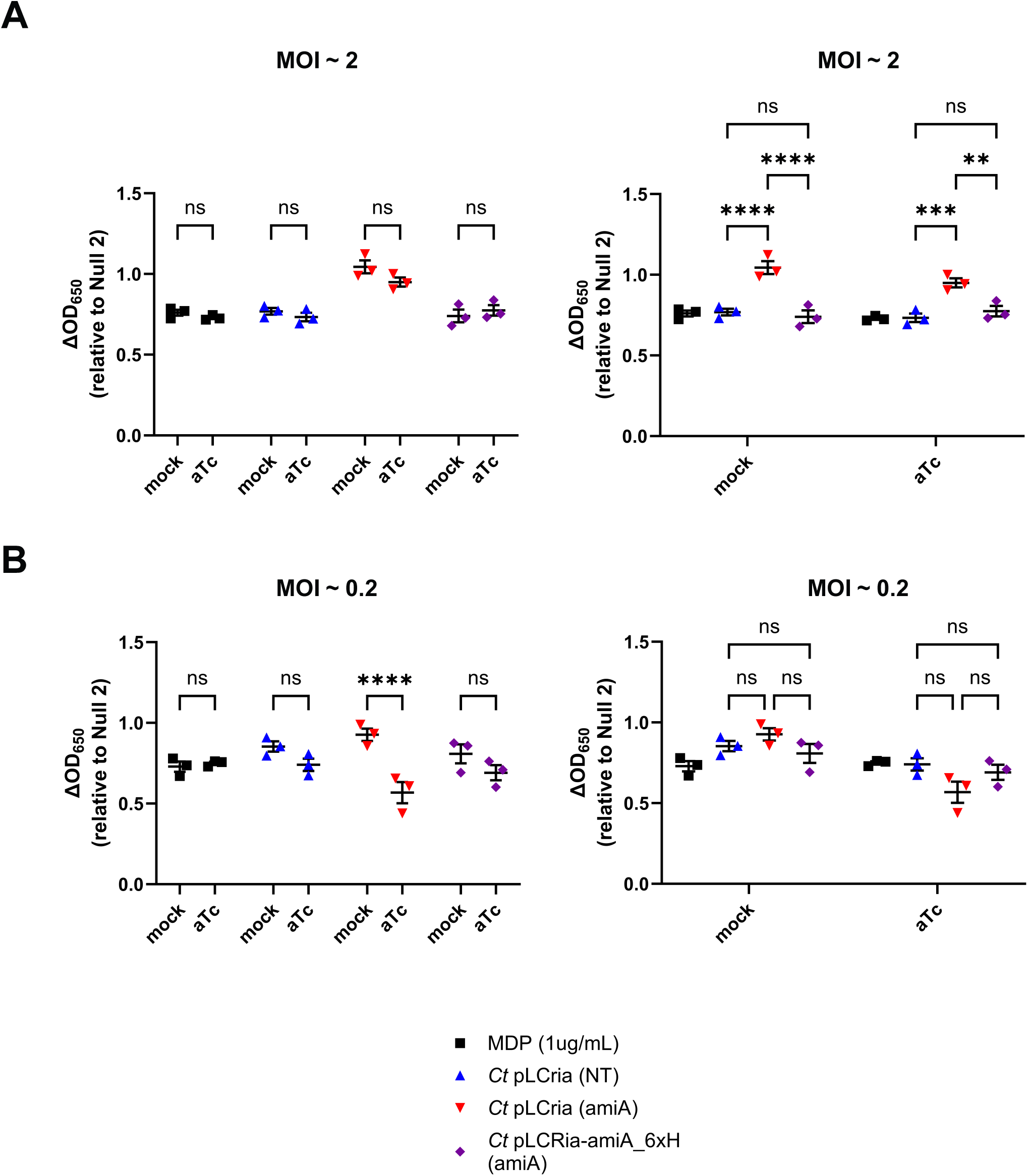
Knockdown of *amiA*_ct_ impacts NOD2 signaling at lower MOIs. hNOD2- expressing HEK293 SEAP reporter cells were infected with *C. trachomatis* strains at MOIs of 2 and 0.2 containing either pLCria (NT), pLCria (*aimA*), or pLCria-amiA_6xHIS (*amiA*). Infections were carried out in the presence / absence of 1nM aTc and supernatants were tested for the presence for SEAP activity 48 hpi. Data points represent three biological replicates, lines indicate the mean of those replicates, and error bars represent standard error of the mean. Groups were compared via two-way ANOVA with multiple comparisons. ****; p ≤ 0.0001, ns, not significant. Data are presented as comparisons between strains **(A)** and between conditions **(B)**.

### Inducers of persistence enhance Chlamydia-specific NOD2 signaling

Under stress-inducing conditions, chlamydial species are capable of entering a state of persistence in which their replicative forms stop dividing, and the developmental cycle will effective pause until the stress-inducing condition is lifted, after which the microbe will resume dividing and transition into its infectious form^65^. There is considerable debate as to whether this aberrant / persistent state is physiologically relevant in the context of active infections^65–71^. We have previously proposed that one of the defining characteristics of this state should be the degree to which it impacts host-microbial interactions at the cellular level^56^. Numerous studies have demonstrated that inducing chlamydial persistence can directly impact the recognition of the organism by various TLRs and NLRs^9,56,72,73^. Given that chlamydia-induced NOD1-signaling has been demonstrated to be impacted in persistently infected cells^56^, we reasoned that NOD2 would be similarly impacted. Upon the addition of D-cycloserine and ampicillin, which respectively inhibit peptidoglycan precursor synthesis^74,75^ and assembly^76^, we found that NOD2 signaling in *C. trachomatis*-infected cells was significantly enhanced **(Fig. 4)**. D-cycloserine was particularly impactful, as NOD2 signaling was observed in DCS-treated, *C. trachomatis*-infected cells as early as 24 hpi and was enhanced as well at later time points **(Fig. 4A)**. When NOD2 signaling was tracked over the span of the chlamydial developmental cycle, signaling from DCS-treated cells began diverging from control and other treatment groups ∼23-24 hpi, while ampicillin-treated cells diverged at ∼38 hpi **(Fig. 4B)**. We also observed enhanced NOD2 signaling in cells that were treated with the iron chelator 2,2′-bipyridyl. Overall, these results demonstrate that induction of aberrance / persistence with the use of PG-targeting antibiotics and iron chelators can significantly impact the degree and kinetics of chlamydia-induced NOD2 signaling.

**Figure 4.**
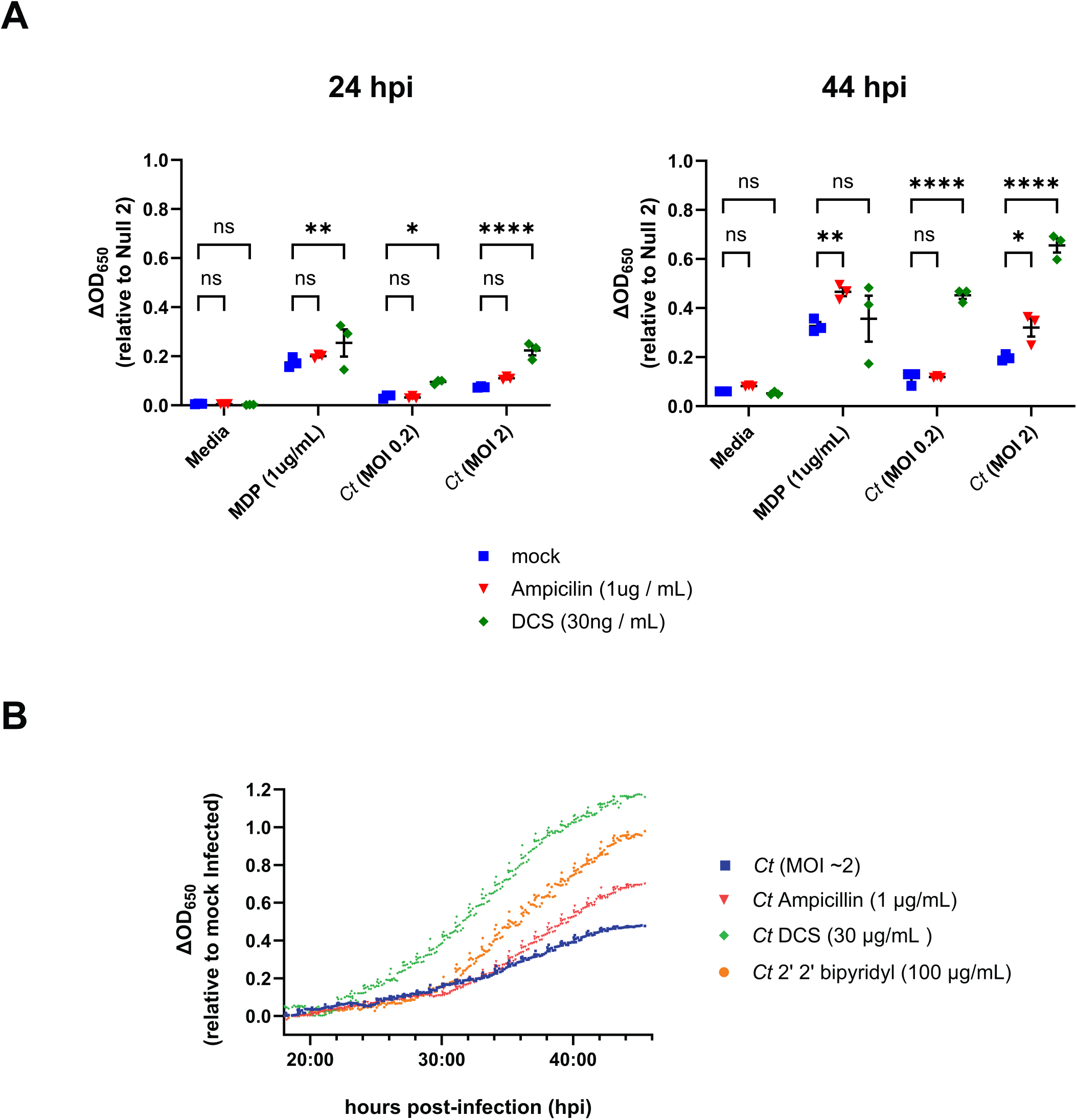
Inducers of persistence enhance Chlamydia-specific NOD2 signaling. **(A)** SEAP activity was measured from the supernatants of hNOD2 and Null2 HEK 293 reporter cells infected with *C. trachomatis* serovar L2 (strain Bu/434) in the presence / absence of antibiotics that target the assembly (ampicillin) and biosynthesis (D- cycloserine; DCS) of peptidoglycan at 24 and 44 hpi. Data points indicate individual biological replicates, lines delineate the mean of those replicates, and error bars represent standard error of the mean. Groups were compared via two-way ANOVA with multiple comparisons. ****; p ≤ 0.0001, **; p≤ 0.01, *; p ≤ 0.05, ns; not significant. **(B)** SEAP activity was measured in the supernatants of infected NOD2-expressing HEK 293 reporter cells grown in HEK-Blue™ Detection media in the presence / absence of peptidoglycan-targeting antibiotics (ampicillin, DCS), and the iron-chelator 2’2 bipyridyl. OD_650_ values for each well were assessed every ten minutes beginning at 18 hpi, and values plotted are relative to mock infected control wells on the same plate. Data is representative of three technical replicates per group.

### Pre-treatment of NOD-expressing cells impacts the development of *C. trachomatis*

Given the difference in signaling kinetics observed between NOD1 and NOD2-expressing HEK293 cells infected with *C. trachomatis* and *C. muridarum*, we wanted to examine whether the resulting signaling cascades adversely impact the growth / development of *C. trachomatis* under native expression conditions. We pre-treated HepG2 cells, which have been previously shown to express both NOD1^77^ and NOD2^78^, with ligands that stimulate NOD1 (L-Ala-γ-D-Glu-mDAP; Tri-DAP) or NOD2 (muramyl dipeptide; MDP) and subsequently infected them with *C. trachomatis* at an MOI of ∼4. At 24 hpi, we then fixed cells and labeled them with antiserum raised against the chlamydial Major Outer Membrane Protein (MOMP). Inclusion numbers were roughly equivalent between all control and test groups examined (data not shown), indicating that NOD-stimulation did not appear to adversely impact inclusion formation. When all MOMP-labeling data was assessed we found cells that were pretreated with MDP prior to infection exhibited smaller inclusion sizes when compared to untreated, Tri-DAP, and MDP + Tri-DAP-treated conditions **(Fig. 5)**. We repeated this assay in two additional cell lines originally derived from cervical (C-33A) and uterine (SiHa) tissue and found that all treatment groups reduced the average inclusion size at 24 hpi.

**Figure 5.**
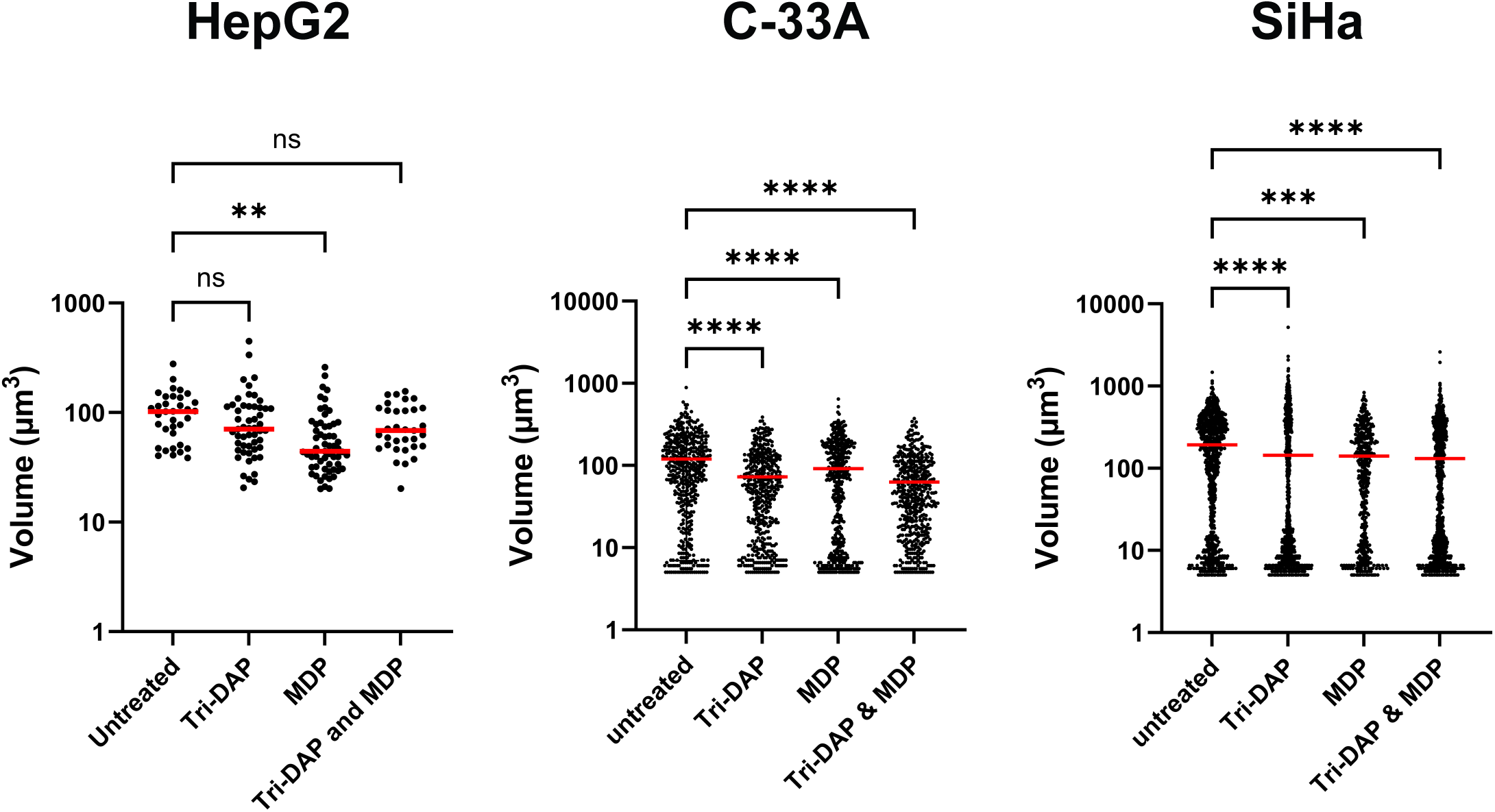
Pre-treatment with NOD1- and NOD2-stimulatory ligands prior to infection with *C. trachomatis* significantly impacts inclusion size in NLR- expressing cell lines. HepG2, C-33A, and SiHa cells were pretreated with Tri-DAP, muramyl dipeptide (MDP), or both, prior to infection with *C. trachomatis* L2 strain Bu/434. At 24 hpi, cells were fixed, inclusions were labeled, counted, and measured. zStacks of MOMP-labeled objects were obtained from a Zeiss 700 confocal microscope and approximate volume measurements were calculated for all inclusions present within 10 imaging fields. Data presented represent all MOMP-labeled objects counted resulting in inclusions > 20 µm^3^; 3 µm across with red lines indicating the mean of each group within a dataset. Significance was assessed via 1-way ANOVA with multiple comparisons. ****; p ≤ 0.0001, ***; p ≤ 0.001, *; p ≤ 0.05, ns; not significant.

We subsequently utilized an inclusion-forming unit (IFU) assay to assess the impact of pre-treatment with NOD1 and NOD2 stimulatory ligands on the chlamydial developmental cycle. We examined the development of *C. trachomatis* L2 strain 434 Bu in HepG2, C-33A, and SiHa cells by assessing the number of viable EBs recoverable at various time points in the pathogen’s developmental cycle. In HepG2 cells, we recovered significantly fewer EBs 48 hours post-infection in the MDP-pretreated group, but these differences were absent when IFUs were recovered at 72 hpi (**Fig. 6A**). We observed similar results in both MDP pretreated C-33A (p ≤ 0.002) and SiHa (p ≤ 0.17) cells **(Fig. 6B,C)** indicating that pre-treatment with PG-derived muropeptides delays chlamydial development in multiple tissue culture cell lines.

**Figure 6.**
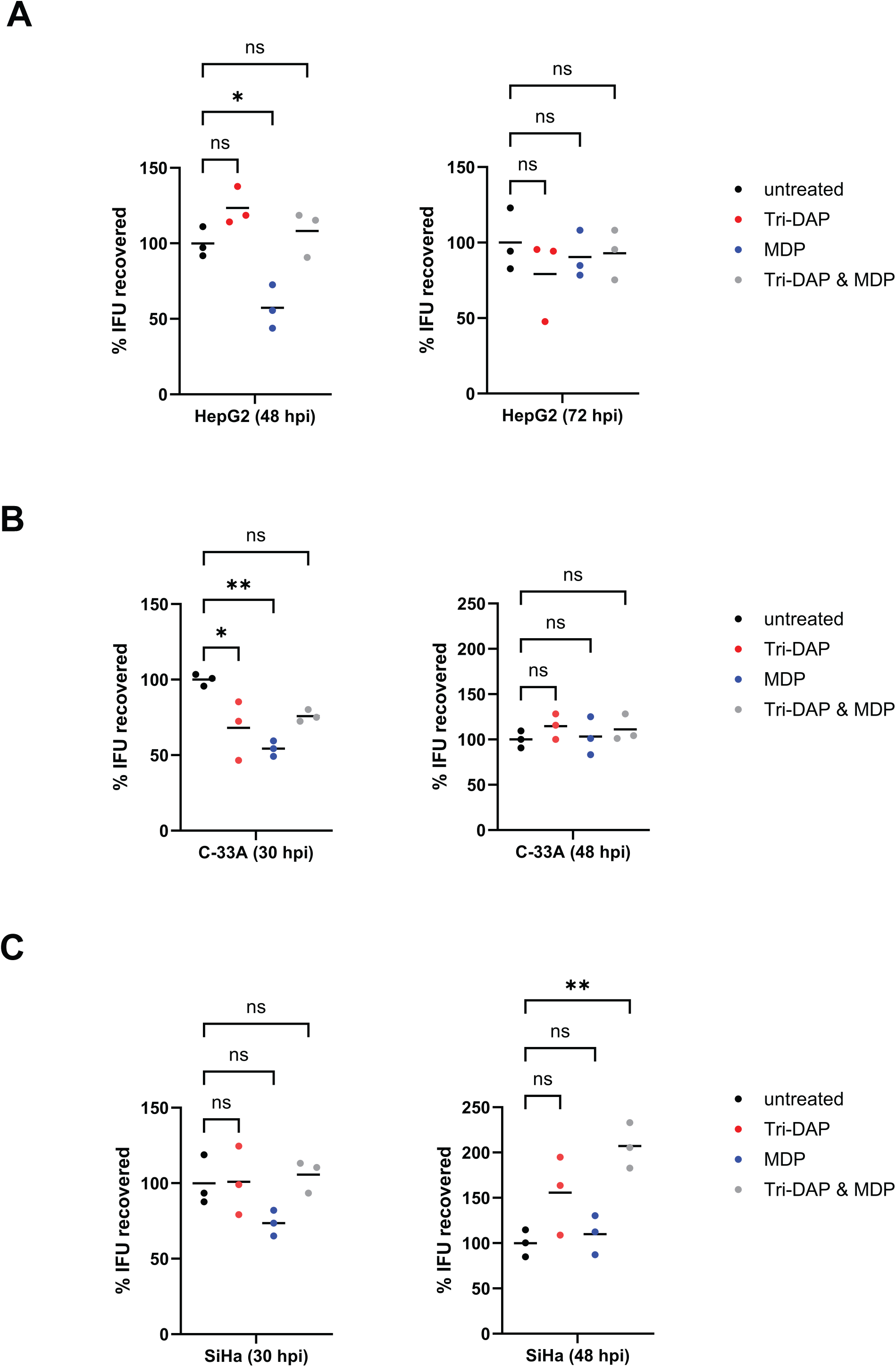
Pre-treatment with NOD2-stimulatory ligands delays the development of *C. trachomatis* in NLR-expressing cell lines. HepG2 **(A)**, C-33A **(B)**, and SiHa **(C)** cells were either left untreated or pre-treated with the NOD1-stimulatory ligand Tri-DAP, the NOD2-stimulatory ligand MDP, or both Tri-DAP and MDP. After 24 hours, cells were infected with *C. trachomatis* and the development of infectious EBs was assessed at the indicated time points (30, 48, and 72 hpi). Lines delineate the mean of 3 separate biological replicates with each replicate displayed as a colored dot. Significance was assessed via 1-way ANOVA with multiple comparisons. **; p ≤ 0.01, *; p ≤ 0.05, ns; not significant.

### Both hNOD2 and mNOD2 recognize C. trachomatis and C. muridarum-specific muropeptides

To conclude this study, we set out to determine whether NOD2 signaling kinetics in Chlamydia-infected cells was unique to i) the human pathogen and/or ii) the human NOD2 receptor. Subsequent experiments were carried out in tandem comparing the NOD2-stimulatory potential of *C. trachomatis* with the murine pathogen *Chlamydia muridarum*. We found that, like *C. trachomatis*, *C. muridarum*-induced hNOD2 expression also appeared to exhibit delayed signaling (**Fig. 7A**). Interestingly, *C. muridarum* also appeared to induce hNOD2 to a greater extent than *C. trachomatis*. We hypothesized that this heightened signaling from *C. muridarum* might be due to one of two factors: differences in the rates of replication between the two pathogens^79^ and / or differences in how each pathogen’s PG was recognized by hNOD2. To test these two non-exclusive possibilities, we replicated our signaling study in HEK 293 cells expressing the murine version of the NOD2 receptor (mNOD2). Both pathogens exhibited similar kinetics in mNOD2 signaling, with SEAP activity only observable at the 44 hpi time point (**Fig. 7B**). Surprisingly, *C. trachomatis* induced higher signaling than *C. muridarum* in mNOD2-expressing cells, potentially indicating that species-specific recognition, rather than differences in bacterial replication rates, is the driving force behind observed differences in NOD2 signaling between these two species.

**Figure 7.**
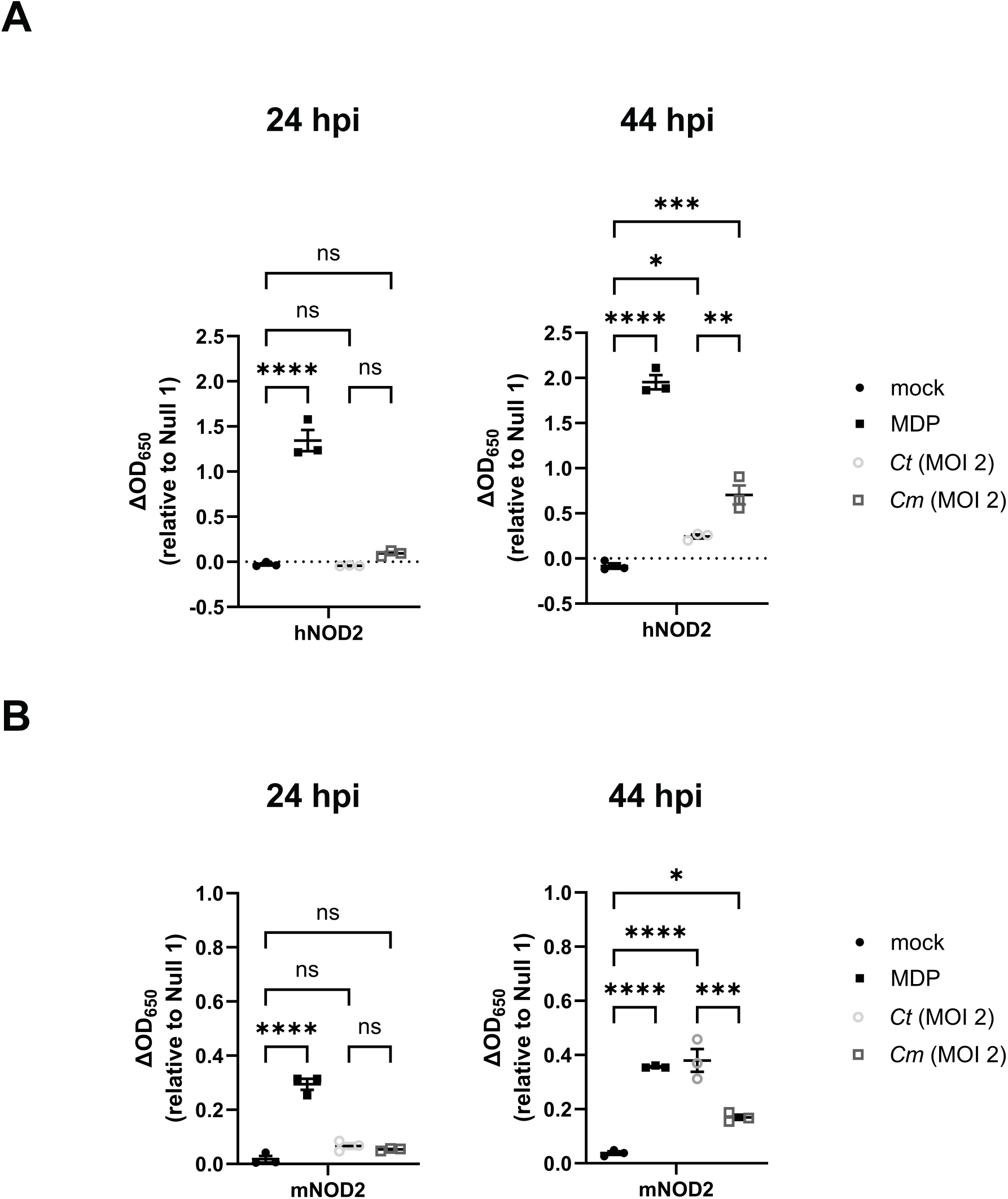
Both hNOD2 and mNOD2 recognize *C. trachomatis* and *C. muridarum*- specific muropeptides. SEAP activity was measured from the supernatants of hNOD2 **(A)**, mNOD2 **(B)**, and Null2 HEK 293 reporter cells infected with *C. trachomatis* serovar L2 (strain Bu/434) at 24 and 44 hpi. Data points represent separate, biological replicates, lines delineate the mean, and error bars represent standard error of the mean. Groups were compared via one-way ANOVA with multiple comparisons. ****; p ≤ 0.0001, ***; p≤ 0.001, **; p≤ 0.01, *; p ≤ 0.05, ns, not significant.

**Figure 8.**
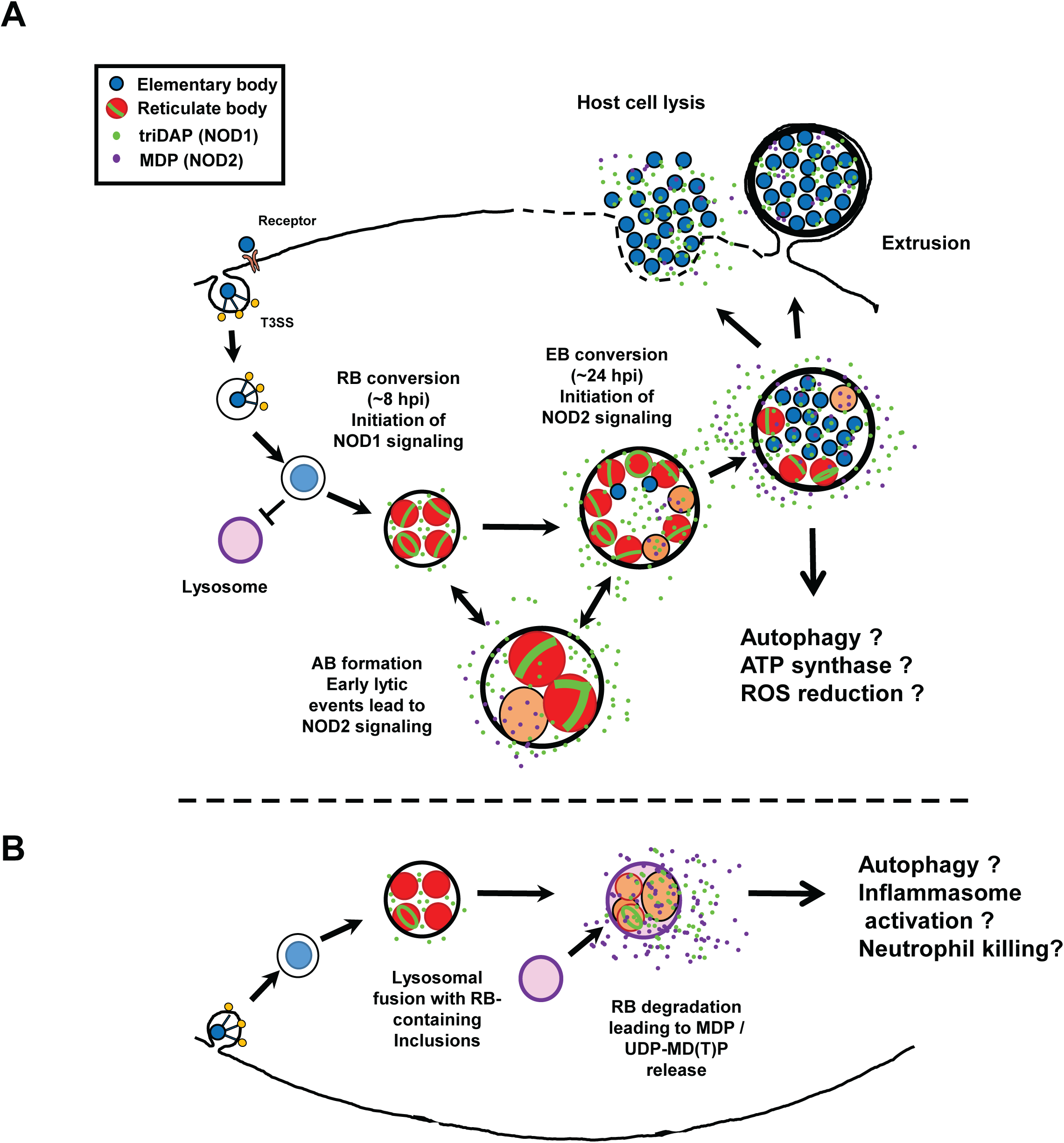
NOD2 signaling as a function of chlamydial lysis during the developmental cycle and ‘persistence’. **(A)** During normal developmental conditions, the chlamydial inclusion avoids fusing with lysosomal compartments in the host cell. After RB conversion, the degradation of PG by AmiA_CT_ during chlamydial replication results in the release of NOD1-stimulatory peptides. Lytic events triggered by developmental form conversion or ‘persistence’ inducers result in the release of partially degraded PG as well as PG precursors, which activate NOD2 signaling. **(B)** In cases were the chlamydial inclusion fuses with lysosomes (such as in phagocytic cells) at later timepoints in the microbe’s developmental cycle (> 8 hpi), we hypothesize that RBs are degraded and NOD2-stimulatory ligands released at a higher abundance, potentially leading to alterations in inflammasome activation.

## DISCUSSION

We have presented data indicating that NOD1 and NOD2 signaling in Chlamydia-infected cells is temporally distinct, and that the kinetics of signaling appear to be impacted by the mechanism(s) the organism utilizes to degrade its PG-derived muropeptides. While these differences appear to be significant, their potential impact on chlamydial development *in vitro* and *in vivo* is less clear. One potential impact of altering NLR signaling might be modulating signaling cascades in such a manner as to be beneficial to the pathogen. While NLRs are associated with upregulation of various cytokine responses^80^, many of which are prevalent during chlamydial infections^81^, one notable difference between NOD1/2 signaling involves a process of direct relevance to most intracellular pathogens: autophagy. NOD2^-/-^ mice exhibit hypersensitivity to viruses as a result of defects in autophagy, specifically mitophagy, resulting in NLRP3 inflammasome activation and the generation of enhanced levels of IL-18^82^. Conversely, chlamydia-infected cells demonstrate the promotion of autophagy^83–85^ and the inhibition of apoptosis^86–89^. While autophagy has been demonstrated to occur in Chlamydia-infected cells in culture, significant levels are generally not observable until later in chlamydial development^83^ and defects in autophagy have been demonstrated to enhance chlamydial growth^84,85^. Given our observation that pre-treatment with NOD2- stimulatory ligands slows chlamydial development in multiple cell lines, we hypothesize that the delayed release of NOD2-stimulatory ligands by *C. trachomatis* represents a pathoadaptation by the microbe, potentially limiting cellular autophagy early in its developmental cycle.

NOD2 is generally thought to enhance cellular autophagy through ATG16L1 in a PI3K, ATG5, and ATG7-dependent manner^90^. ATG16L1 has been shown to restrict the expansion of the chlamydial inclusion, but is re-directed by the secreted chlamydial protein CT622/TaiP to enable vesicular traffic to the inclusion^91^. Given that we see a reduction in inclusion size in MDP pre-treated cells **(Fig. 5)**, it is tempting to speculate that NOD2 signaling impacts chlamydial development by recruiting the majority of cellular ATG16L1 towards autophagy, thus leaving less free to be utilized by TaiP to recruit vesicular traffic to the growing chlamydial inclusion. The regulation of NOD2- stimulatory ligands by chlamydial species may also exhibit tissue-specific effects, as MDP has recently been shown to increase oxidative respiration and ATP production while decreasing oxidative stress in human intestinal epithelial cells^92^. Experiments investigating these interesting possibilities are currently ongoing.

Many of our observations in this study mirrored those previously reported regarding *Chlamydia*-induced TLR9 signaling^9^. Both *Chlamydia*-induced NOD2 and TLR9 signaling occur late in infection, can be significantly reduced if protein synthesis is inhibited prior to phase conversion, and are significantly enhanced when infected cells are treated with inhibitors of either PG or LOS biosynthesis. Given that PG and LOS inhibitors are known to interrupt phase conversion and impact the integrity of the bacterial membrane, we hypothesize that their presence results in an increase in chlamydial lytic events, resulting in PG and gDNA release. The observation that *C. muridarum* and *C. trachomatis* appear to differ in the intensity of their signaling in a host-specific manner is another notable finding. While both hNOD2 and mNOD2 recognize MDP and both have similar downstream signaling pathways, some differences have been observed such as the degree to which mutant alleles are capable of suppressing IL-10 expression^93^. Species-specific differences have been noted for other TLRs and NLRs with regard to their regulation, ligand specificity and function^94^, and differences in ligand specificity and sensitivity have been previously demonstrated for both hNOD1 and mNOD1^95^.

As persistence in *Chlamydia* species often leads to a disruption in membrane integrity^96^, we were not surprised by the finding that NOD2 levels increased significantly as a result of persistence induction. Our observation of enhanced NOD2 signaling in cells that were treated with the iron chelator 2,2′-bipyridyl was unexpected, as this condition has previously been demonstrated to significantly reduce chlamydia-specific NOD1 signaling at early infection time points^56^. The finding that DCS substantially impacts not only the degree but also the timing of Chlamydia-specific NOD2 signaling was particularly interesting. This observation matches those previously made investigating Chlamydia-derived, NOD2-stimulatory ligands present in HeLa cell lysates^57^, indicating that the enhancement is not simply a result of the sonication steps involved in preparing the material for evaluation. DCS is a D-alanine analog, originally isolated from *Streptomyces*^97,98^, and historically its mechanism of action has been understood to be the result of it competitively binding to D-alanine racemase; Alr^75^ and D-alanine-D-alanine ligase; Ddl^99^. Chlamydia species lack canonical Alr and DadX homologs, but encode GlyA enzymes capable of D-Ala racemase activity, which is also inhibited by DCS^100^. The inactivation of these two enzymes should inhibit PG biosynthesis. As chlamydial species do not encode MpaA homologs that would normally function to hydrolyze γ-D-glutamyl-diaminopimelic acid^101^, the result of treatment should be the accumulation of UDP-MurNAc-L-Ala-D-Glu-*meso*-DAP (UDP-MTP) in the chlamydial cytoplasm. While muramyl tripeptide (MTP) is not well recognized by NOD2^95^, the presence of a Uridine Diphosphate (UDP) carrier group in the PG precursor molecule (UDP-MTP) substantially enhances its recognition by NOD2 receptors^102^. This would support the hypothesis that NOD2 signaling from DCS-treated Chlamydia-infected cells is likely due to the release of PG-precursors (namely UDP- MTP) by aberrant RBs. Given that UDP-MTP is normally localized to the bacterial cytoplasm, this would appear to indicate that chlamydial RBs under these conditions are undergoing appreciable levels of cellular lysis. Alternatively, it has been recently suggested that DCS is capable of being utilized as a substrate by both L,D- and D,D-transpeptidases and is capable of being incorporated into the stem peptide of PG at the 4^th^ and 5^th^ position, respectively^103^. It is presently unclear whether these DCS-containing peptide stems are capable of being crosslinked, or whether their presence significantly impacts immunorecognition by NLRs.

The chlamydial amidase AmiA_CT_ appears to meaningfully impact Chlamydia-induced NOD2 signaling (**Fig 3**). During PG turnover, AmiA_CT_ cleaves stem peptides from MurNAc^61^, and only once this has occurred can lytic transglycosylases (ie. SpoIID_CT_) bind to and subsequently degrade the newly denuded PG glycan stands. However, we have also shown that disruptions in chlamydial membrane integrity, alterations in PG biosynthesis, and chlamydial lytic events that can occur during phase conversion can all influence chlamydial-derived NOD2 signaling. The enhanced background signaling and slightly enlarged RBs present in our uninduced, AmiA_CT_ knockdown strain would appear to indicate low levels of dCAS9 expression. When baseline NOD2 levels are equivalent between all three strains when using a lower MOI, we see a significant decrease in NOD2 signaling upon induction specific to our knockdown strain **(Fig. 3B)**. Assuming that SpoIID_CT_ is unable to process glycan strands prior to amidase activity, it is somewhat puzzling that the knockdown of amidase activity would impact NOD2 activity. As we observe elevated NOD2 signaling at higher MOIS and lower activity under induction conditions at lower MOIs, we speculate that this data may represent different origins of NOD2-stimulatory ligands and potentially support the hypothesis that PG fragments are not the principle NOD2-stimulatory ligands being released by *C. trachomatis*. Given our DCS-treatment data (**Fig. 4**), we conclude that PG-precursors that still retain their UDP carrier motif likely make up a substantial portion of the NOD2-stimulatory ligands present in Chlamydia-infected cells, though subsequent experimentation will be needed to further validate this.

While this study effectively demonstrates potential differences in the temporal kinetics of Chlamydia-induced NOD1 and NOD2 signaling, it is important to point out that these phenotypes will almost certainly differ in hematopoietic vs. non-hematopoietic cell types. While *C. trachomatis* may effectively delay NOD2 signaling until late in development by cleaving stem peptides during the natural PG degradative processes associated with its division cycle, this may not significantly impact the production of NOD2-stimulatory PG fragments generated by cellular responses to infection, such as the potential degradation of RBs within lysosomal compartments within hematopoietic cells **(Fig. 7)**. However, as only chlamydial RBs synthesize and maintain PG^13,15^, its availability for NOD1/2 interactions in immune cells would largely depend on the kinetics of the pathogen trafficking to lysosomes. Given that the EB-to-RB transition occurs roughly 8 to 12 hpi, this is presumably plenty of time for EB-containing vesicles to traffic to lysosomes^104,105^. It is tempting to speculate that the origins of EB-to-RB transition kinetics may have arisen, in part, as a means of dampening potential immune responses generated in hematopoietic cells that are incompatible with chlamydial replication and development. Alternatively, as the major NOD2-stimulatory ligand (MDP) is a known activator of NALP3/NLRP3^106^, its release during RB degradation could contribute to neutrophil killing via NLRP3 inflammasome^107^. Work investigating how the timing of inclusion-lysosomal fusion impacts these downstream processes in a cell-type specific manner is currently under active investigation.

## CONCLUSIONS

In this study, we demonstrate that chlamydia infection can be detected by both NOD1 and NOD2 receptors *in vitro*, but that the kinetics of that signaling differs significantly with NOD2 signaling lagging behind that of NOD1. Additionally, we effectively demonstrate that the timing and intensity of NOD2 signaling can be impacted by a variety of factors: chlamydial / host species, incidence of chlamydial lysis, the relative activity of the chlamydial amidase enzyme, and the induction of aberrance / ‘persistence’ phenotypes. Our results strongly suggest that chlamydial PG precursors, rather than degraded PG fragments, are the primary stimulators of NOD2. Given our observations that pre-stimulating host cells with MDP delays chlamydial development and NOD2’s unique association with cellular autophagy induction, we reason that the delay in NOD2-induction by chlamydial species directly benefits the development of these obligate, intracellular pathogens.

## MATERIALS AND METHODS

### Reagents

Anti-MOMP antisera (LS-C123239) was purchased from LSBio. LpxC inhibitor LPC-011 was graciously provided by Dr. Pei Zhou (Duke University). Muramyl dipeptide (MDP) and L-Ala-γ-D-Glu-mDAP (Tri-DAP) were purchased from InvivoGen.

### Bacterial Strains and Cell Lines

*C. trachomatis* serovar L2 strain 434/Bu, *C. muridarum* strain Nigg, and *E. coli* strain MG1655 were provided by Anthony Maurelli (University of Florida). *C. trachomatis* and *C. muridarum* stocks were generated utilizing HeLa-USU cells (also provided by Anthony Maurelli) unless otherwise noted. *C. trachomatis* strains -pL2 transformed with pLCria (NT), pLCria (*amiA*), and pLCria-amiA_6xHIS (amiA) vectors were provided by Scot Ouellette (University of Nebraska Medical Center) and are characterized in Dannenberg *et al*. (manuscript in prep). Whole cell lysate (‘crude’) freezer stocks of *C. trachomatis* and *C. muridarum* EBs were generated from HeLa cells 40 hours post infection and stored at -80° C in sucrose phosphate glutamic acid buffer (7.5% w/v sucrose, 17 mM Na_2_HPO_4_, 3 mM NaH_2_PO_4_, 5 mM L-glutamic acid, pH 7.4) until use. Stocks were titered via inclusion forming unit (IFU) assay (described below). HEK-Blue-hNOD1, -hNOD2, -mNOD2, and -Null1 cells were purchased from InvivoGen and propagated according to the manufacturer’s instructions. Cell lines were passaged in high-glucose Dulbecco’s modified Eagle medium (DMEM; Gibco) and 10% fetal bovine serum (FBS; HyClone). HepG2 cells were purchased from ATCC and propagated according to their instructions. All cell lines were checked for mycoplasma contamination 2 passages after the initial liquid nitrogen thaw, and every subsequent 10 passages.

### HEK-Blue hNOD2, mNOD2, Null1 and Null2 NF-κB reporter assay

HEK-Blue cells expressing human or murine NOD2 and carrying the NF-κB SEAP (secreted embryonic alkaline phosphatase) reporter gene (InvivoGen) were used according to the manufacturer’s instructions and adapted to assess NLR-specific NF-κB activity induced via live *C. trachomatis* and *C. muridarum*. Briefly, 3 × 10^5^ cells/ml were plated in 96-well plates (total reaction volume of 200 μl per well [∼6.0 × 10^4^ cells per well]) and allowed to settle/adhere overnight at 37°C. Media was then removed and replaced with 200 µl of medium containing either *C. trachomatis* or *C. muridarum* (at the MOIs indicated) or known NOD-stimulatory ligands (triDAP and MDP). For experiments examining the impact of conditionally enhancing the developmental cycle of *Chlamydia* spp. on NOD signaling, multiple MOIs were used. Plates were then centrifuged for 1 h at 2,000 × *g* and subsequently incubated in a CO_2_ incubator at 37°C. Cell supernatants were collected at indicated time points for subsequent analysis of SEAP activity. A colorimetric reporter assay was then utilized to quantify the abundance of SEAP in cell supernatants. Twenty microliters of supernatant collected from infected cells was added to 180 μl of the SEAP detection solution (InvivoGen), followed by incubation at 37°C for ∼6 h. SEAP enzymatic activity was then quantified using a plate reader set to 650 nm. Infected cells were compared to uninfected cells (negative control) and cells treated with triDAP / MDP (positive controls). To ensure that changes in alkaline phosphatase activity were NLR-dependent under each of the experimental conditions tested, all experiments were carried out in parallel in either HEK-Blue-Null1 or HEK-Blue-Null2 cell lines, which contains empty expression vectors but lack NOD1/2. Each HEK-Blue SEAP reporter assays was always carried out in three separate experiments, statistical analysis was conducted by either 1- or 2-way ANOVA, and significance values were analyzed by utilizing Sidak’s multiple-comparison test. For real-time tracking of NOD2 signaling, hNOD2 reporter cells were plated, and infected as described above, but instead of DMEM / 10% FBS, cells were incubated in HEK-Blue Detection cell culture medium. Plates were placed in a Stratus microplate reader (Cerillo), which itself was placed in a CO_2_ incubator at 37°C. OD650 readouts were obtained beginning at 18hpi every 10 minutes for the subsequent 28 hours.

### *C. trachomatis* Infections of HepG2, C-33A, and SiHa cells

∼2.5 x 10^5^ cells / mL were spun down and resuspended in infection medium containing ∼1x10^6^ IFU of *C. trachomatis* (MOI ∼4). A MOI of 4 was used in order to maximize the number of inclusions visible per field of view. 2.5 x 10^5^ cells were then added to each well of a 24 well plate and placed in the incubator for 24 or 44 hours. For pre-stimulation experiments, ∼2.5 x 10^5^ cells / mL were spun down and resuspended in infection medium + / - 20 µg/mL triDAP, 20 µg/mL muramyl dipeptide (MDP), or 10 ug/mL of each and incubated overnight at 37° C. The next morning, cells were spun down and resuspended in infection medium containing ∼1x10^6^ IFU of *C. trachomatis* (MOI ∼4), 2.5 x 10^5^ cells were then added to each well of a 24 well plate and placed in the incubator (at 37° C). Cells were then either fixed for inclusion visualization or lysed for the enumeration of viable EBs via IFU assay.

### Fluorescence Microscopy / MOMP-labeling

Briefly, HeLa cells were infected with *C. trachomatis* L2 434/Bu as described above. A MOI of 4 was used, in order to ensure a higher number of visible MOMP-containing objects within a single imaging field. At indicated time points infection medium was removed, and cells were washed three times with 1× PBS. Cells were fixed and permeabilized with methanol for 5 min and gently washed three times with 1× PBS. Cells were then further permeabilized and blocked as described above. Cells were then incubated with anti-MOMP (1:500) for 1 hour at room temperature, followed by several washes in 3% BSA, and subsequent incubation with anti-goat conjugated antibodies (1:1000 Alexa594). For visualizing cell nuclei, cells were incubated with Hoechst stain for ∼10 min and then washed with 3% BSA and 1× PBS. Coverslips were mounted on slides with ProLong gold antifade mounting medium and stored in the dark at 4°C prior to imaging via structured- illumination (Elyra PS.1) microscopy. For inclusion size measurements, imaging was conducted on a Zeiss 980 confocal microscope. zStacks were acquired at random for ∼10 fields of view per condition, and inclusion volume dimensions were calculated utilizing the 3D object counter addon (ImagJ / Fiji).

### Inclusion-forming Unit (IFU) Assay

Viable EBs were assessed as previously described^9^. Briefly, 96-well tissue culture-treated plates were seeded 24 hours prior to infection with ∼40,000 cells per well. On the day of infections, bacterial suspensions from crude cell lysates were thawed on ice, and serially diluted in DMEM / 10% FBS + 0.5 ug/mL cycloheximide. Spent media was removed from each well and 200 µl of each chlamydia dilution was added to each well. Plates were then centrifuged at 3000 x g at 35° C for 1 hour, then incubated at 37° C with 5% CO2 for an additional 23 hours. Cells were then fixed with ice cold methanol for 10 minutes, washed with 3 x PBS, blocked with 3% BSA for 1 hour, incubated with goat anti-MOMP (LSBio) for 1 hour at RT, washed 3 x with PBS, incubated with donkey anti-goat Alexa Fluor 488 (Invivogen) for 1 hour at RT, and washed 3 x with PBS. Plates were then examined under an epifluorescence microscope (Olympus) within 24 hours. Total inclusion forming units (IFU) were calculated by counting the number of visible inclusions for 10 different imaging fields (per condition) using the 40x objective.

## Data Availability

All data generated in this manuscript will be made available without restriction upon request. This includes all super resolution imaging data, all reporter cell line SEAP data, and chlamydial IFU calculations.

## ACKNOWLEDGEMENTS

We would like to thank Dr. Anthony Maurelli (University of Florida) and his laboratory for providing us with the strains used in this work as well as helpful feedback during the development of this project. We would like to thank Dr. Scot Ouellette (UNMC) for providing us with strains for conducting the AmiA_CT_ knockdown experiments. We would also like to thank Dr. Pei Zhou (Duke University) for graciously providing us with the LpxC inhibitor used in this work. This work was supported by a MIRA ESI award (R35 GM138202) and a USU faculty start up award (HP73LIEC18) to GL. Funding was also provided by the Deutsche Forschungsgemeinschaft (DFG, German Research Foundation), project-ID 398967434 —TRR261 (BH) and 390536577 (BH), and the funding scheme FEMHABIL, Medical Faculty, University of Bonn (BH). The funders had no role in study design, data collection and interpretation, or the decision to submit the work for publication. The opinions and assertions expressed herein are those of the author(s) and do not necessarily reflect the official policy or position of the Uniformed Services University or the Department of Defense.

## AUTHOR CONTRIBUTIONS

Conceptualization and Design, IL, BH, GL; Data Curation and Formal analysis, GO, IL, ZW, SD, AD, and GL; Investigation, Methodology, Validation, Visualization, IL, AD, GO, ZW, JB, and GL, Writing – original draft, GL; Writing – review & editing, GO, IL, ZW, SD, JB, BH, and GL; Funding acquisition, Project administration and Supervision, BH and GL. All authors read and approved the final manuscript.

